# A Chlorinated Diketopiperazine Antibiotic Targets *Mycobacterium Tuberculosis* DNA Gyrase

**DOI:** 10.1101/2025.03.10.642354

**Authors:** Libang Liang, Jeffery Quigley, Monique Theriault, Akira Iinishi, Rachel Bargabos, Madeleine Morrissette, Megan Ghiglieri, Tom Curtis, Rachel Corsetti, Sangkeun Son, Bishwarup Sarkar, Kim Lewis

**Author notes:** Corresponding Author: Kim Lewis. These authors contributed equally (Libang Liang, Jeffery Quigley). The authors declare no competing financial interest.

## Abstract

We describe a novel macrocyclic peptide, speirobactin, produced by *Photorhabdus* that selectively kills *Mycobacterium tuberculosis*. A non-ribosomal peptide synthase (NRPS) containing two linear modules codes for the synthesis of speirobactin. The biosynthetic operon contains a pentapeptide-repeat protein as a resistance gene. Genomic analysis of speirobactin-resistant mutants of *M. tuberculosis* led to identification of DNA gyrase as the molecular target. The mutations were recreated by allelic replacement and show that DNA gyrase is the only target. Transcriptome analysis of *M. tuberculosis* treated with antibiotics shows that speirobactin clusters close to fluoroquinolones, supporting its action against the DNA gyrase.

## Introduction

*Mycobacterium tuberculosis* is the leading cause of death from a single infectious agent according to the World Health Organidzation (https://www.who.int/news-room/fact-sheets/detail/tuberculosis). The standard of care for treating drug susceptible *M. tuberculosis* requires an intensive treatment with four antibiotics -rifampin, isoniazid, pyrazinamide, and ethambutol for 8 weeks, followed by 18 weeks of extended treatment using isoniazid and rifampin. In 2021, results of a phase III clinical trial supported a 4 months treatment using rifampentine and moxifloxacin was noninferior to the standard 6 month treatment (1). Following this study, the Centers for Disease Control and Prevention issued an interim guidance to recommend the 4-month rifampentine-moxifloxacin regimen for treating patient aged over 12 with drug-susceptible pulmonary tuberculosis. The prolonged treatment and patient incompliance have given rise to drug-resistant *M. tuberculosis* strains (2, 3). New antibiotics are needed to combat drug-resistant *M. tuberculosis*.

Streptomycin was the first antibiotic able to treat tuberculosis. Another natural product compound, rifampicin (or rifabutin), is still an important component of the treatment regimen. Yet, resistance to the first line antibiotics such as rifampicin is a major concern for tuberculosis. In search of additional compounds to treat tuberculosis, we introduced differential screening and focused on untapped sources of secondary metabolites. A very large background of toxic and known compounds is a formidable bottleneck for antibiotic discovery. Screening in parallel against *M. tuberculosis* and a different species such as *Staphylococcus aureus* resolves this bottleneck. Only compounds acting selectively against *M. tuberculosis* are considered, and with selective activities these compounds cannot be generally toxic. Selectivity against a particular species of bacteria also suggests lack of action against eukaryotic cells. This approach coupled with a screen of uncultured bacteria, has led to the discovery of lassomycin, an inhibitor of the mycobacterial ClipP1P2C1 protease from *Lentzea kentuckyensis* sp., amycobactin, an inhibitor of mycobacterial SecY protein exporter produced by *Amycolatopsis* sp., and evybactin a DNA gyrase inhibitor produced by *Photorhabdus* sp. smuggled into cells by the transporter BacA (4– 6). Searching for untapped sources of novel compounds, we have been screening *Photorhabdus*, symbionts of nematode gut microbiome. The entomopathogenic nematodes infect insect larvae and release *Photorhabdus* that make antimicrobial compounds to fend off competitors. These compounds should be non-toxic to the nematode. This approach led to the discovery of darobactins and dynobactins, inhibitors of the BamA outer membrane chaperone of Gram-negative bacteria (7, 8). In this study, a differential screen of *Photorhabdus* resulted in the discovery of speirobactin, an inhibitor of DNA gyrase that acts selectively against *M. tuberculosis*.

## Results and Discussion

### Bioassay Guided Isolation and Characterization of Speirobactin

A small set of 17 *Photorhabdus* isolates (Table S1) were fermented in 15 different media, and supernatants were withdrawn over time. This resulted in 765 samples which were centrifuged to remove cellular components. The supernatants were tested against *M. tuberculosis* in microtiter plates and counter-screened against *S. aureus* in an agar diffusion assay. To facilitate the screen, we used a BSL2 strain of auxotrophic *M. tuberculosis* mc2 expressing mCherry, allowing to detect growth by measuring fluorescence. The centrifuged *Photorhabdus* supernatant produced 10, 8, and 8 hits against *M. tuberculosis* with over 90% inhibition on day 2, day 5, and day 8 respectively. After consolidation, they came from three producer strains (*P. temperata* He86, *P. temperata* K122, and *P. temperata* KLE11398) fermented in different media (Table S2). In the counter-screen against *S. aureus*, the crude extracts from *P. temperata* KLE11398 with hits against *M. tuberculosis* showed strong inhibition zones with *S. aureus* while extracts from the other two producers had mild inhibition. Therefore, *P. temperata* He86 and *P. temperata* K122 extracts were followed up since the inhibition against *S. aureus* was relatively weak. One fermentation medium (Table S3) was chosen for each of the two producers for further scale-up.

The two producers were scaled up by liquid fermentation (*P. temperata* He86 in medium M15 and *P. temperata* K122 in Medium M4) for five days, followed by ethyl acetate extraction. The crude extracts were fractionated by C18 column chromatography and tested against *M. tuberculosis*. The active fractions were further fractionated by High Performance Liquid Chromatography (HPLC) until the active fractions contained only one compound based on HPLC-UV.

The purified samples were analyzed by Liquid Chromatography - High Resolution Mass Spectrometry (LC-HRMS) to identify the exact mass under electrospray ionization (ESI) in positive mode. Compounds from both *P. temperata* He86 and *P. temperata* K122 shared the same HPLC retention time and the same exact mass with an *m/z* [M+H]^+^ of 376.0801, suggesting the same active compound was produced by both strains. There was a characteristic isotopic ion with an *m/z* of 378. 0772 at about a third of the peak intensity, suggesting the molecule contained one chlorine. With one chlorine in the molecule, there was only one molecular formula C_16_H_14_N_5_O_4_Cl under 5 ppm mass deviation. Therefore, the molecular formula C_16_H_14_N_5_O_4_Cl was assigned for the neutral molecule, which we named speirobactin. Another 10 L scale up using the producer *P. temperata* He86 followed by extraction and purification afforded 3 mg of the purified speirobactin for structure elucidation.

1D and 2D NMR analysis (^1^H, ^13^C, COSY, ^1^H-^13^C HSQC, ^1^H-^13^C HMBC, TOCSY, NOESY) established the diketopiperazine backbone structure of speirobactin consisting of a tryptophan moiety and an asparagine moiety (Fig. S1-Fig. S7, Table S4). The amine proton H-15 has HMBC correlations to C-16, C-13, and C-21, and the aromatic proton H-14 has HMBC correlations to C-21, C-13, and C-16, establishing the five-member ring in the tryptophan moiety. Aromatic protons H-17 and H-20 showed HMBC correlations to C-21, and H-17 showed the HMBC correlation to C-16, suggesting the three aromatic protons H-17, H-19, and H-20 were in an aromatic system connected to the five-member ring. A COSY correlation between H-19 and H-20 suggested C-19 was connected to C-20. H-17, H-19 and H-20 showed HMBC correlations to C-18, the last carbon in the six-member ring of tryptophan. Although H-19 showed an HMBC correlation to C-17, there was lack of TOCSY correlations between H-17 and H-19, which indicated C-18 was attached to a heteroatom. The carbon chemical shift of C-18 could not support an oxygen or nitrogen substitution; based on the molecular formula, a chlorine was assigned at the C-18 position. H-12 showed an HMBC correlation to C-14, and H-12 and H-14 had an NOE correlation, establishing the H-12 as the β-proton in the tryptophan. The presence of the strong NOE correlation between H-12 and H-14 and the absence of NOE correction between the NOE correlation between H-12 and NH-8 established the Z configuration of the double bond. Additionally, the Z configuration was deduced by the downfield proton chemical of H-12 (*δ*_H_ 6.92) shielded by the carbonyl at C-10, which had been reported in multiple structurally related diketopiperazines (9, 10). H-12 also showed HMBC correlations to C-9 and the carbonyl carbon C-10, suggesting C-9 was the α-carbon of the tryptophan. A double bond was assigned between C-9 and C-12 based on their chemical shifts and only one proton attached to C-12.

Another amide proton H-11 showed HMBC correlations to C-10, C-9 and the asparagine carbonyl C-7. COSY correlations between H-11 and H-6 as well as between H-5 and H-6 suggested H-5, H-6, and H-11 were in the same spin system. H-5 as the β-proton showed an HMBC correlation to the carbonyl C-7, and another carbonyl C-4. An NOE correlation between H-5 and the amide proton H-3 established the assignment of the asparagine moiety. A carbon signal at 154.1 ppm was observed in the ^13^C spectrum, which was the last unassigned carbon from the molecular formula. Together with the last unassigned nitrogen, a carbamide moiety was proposed to attach to the side chain of the asparagine. MS/MS analysis of the speirobactin [M+H]^+^ ion 376.08 resulted in ions with m/z of 359.04, 333.15, and 316.06, which corresponded to loss of NH_2_, CONH_2_, and NH_2_CONH_2_ from the asparagine side chain (Fig. S8), confirming the carbamide assignment. Finally, the connection between the carbonyl C-7 and the tryptophan N-terminal was assigned to complete the degree of unsaturation and forming a diketopiperazine core structure.

We noticed the structure of speirobactin contained a chiral center at the asparagine moiety. The stereochemistry was analyzed with a modified Marfey’s method. The analysis showed the hydrolyzed speirobactin contained both D- and L-aspartic acid (Fig. S10). Further, the racemic mixture of speirobactin was separated using chiral HPLC (Fig. S9) and the MIC for (6R)-speirobactin was 0.25 µg/mL while (6S)-speirobactin was 4 µg/mL. We also acknowledge that the activity from (6S)-speirobactin could come from the contamination of (6R)-speirobactin.

The speirobactin MIC against *M. tuberculosis* was 0.25 µg/mL, and 16 µg/mL against the counter-screen organism *S. aureus*. This shows that speirobactin is a highly potent and selective antibiotic acting against *M. tuberculosis*. Speirobactin had low activity against human pathogens and gut commensals, confirming its selectivity against *M. tuberculosis* (Table 1). Notably, while speirobactin was inactive against wild type *E. coli*, it was highly potent against *E. coli* WO153, a strain with a defective permeability barrier, deficient in production of lipopolysaccharide and lacking TolC, a porin for docking MDR pumps (11). As expected for a selective compound, speirobactin is non-toxic to the tested mammalian cells (Table 1).

**Table 1.** Activity of speirobactin against pathogens, commensals and human cells.

| Organism and genotype | Speirobactin concentration (µg/mL) |
| --- | --- |
| Pathogenic bacteria (MIC) |  |
| *Mycobacterium tuberculosis* H37Rv | 0.25 |
| *Mycobacterium smegmatis* mc<sup>2</sup>155 | 16 |
| *Borrelia burgdorferi* BbP1286 | 1.25 |
| *Staphylococcus aureus* HG003 | 16 |
| *Pseudomonas aeruginosa* PAO1 | 16 |
| *Klebsiella pneumoniae* ATCC 700603 | >128 |
| *Escherichia coli* MG1655 | >128 |
| *Escherichia coli* WO153 | 0.25 |
| *Acinetobacter baumannii* ATCC17978 | >128 |
| Human anaerobic commensals (MIC) |  |
| *Bacteroides fragilis* KLE 2244 <sup>a,b</sup> | 8 |
| *Bacteroides stercoris* KLE 2537<sup>a,b</sup> | 16 |
| *Veillonella ratti* KLE 2365 <sup>a,b</sup> | 32 |
| *Lactobacillus reuteri* LTH5448 <sup>a</sup> | >128 |
| *Clostridium perfringens* KLE 2523 <sup>a,b</sup> | 32 |
| *Clostridium tertium* KLE 2303 <sup>a,b</sup> | 8 |
| *Enterococcus faecalis* KLE 2341 <sup>a,b</sup> | 4 |

| Cell lines (IC<sub>50</sub>) |  |
| --- | --- |
| Hek293 | >128 |
| Fadu | >128 |
| HepG2 | >128 |

- a. Cultured under anaerobic conditions - b. Human stool isolates, KLE (Kim Lewis laboratory collection)

### Biosynthetic Gene Cluster of Speirobactin

Genome sequencing and assembly of the producers *P. temperata* He86 and *P. temperata* K122 resulted in two genomes both at 5.6 M bp. AntiSMASH 7.0 (12) was used to profile biosynthetic gene clusters (BGCs) in the two genomes, and 26 and 22 BGCs were identified for *P. temperata* He86 and *P. temperata* K122, respectively. Based on the elucidated structure of speirobactin, its chemical backbone contains tryptophan and asparagine. We started to focus on nonribosomal peptide synthetases (NRPSs) and narrowed down to specific pathways that incorporate two amino acids. In this way, the expected NRPS should contain two linear modules. All BGCs containing NRPSsfrom the AntiSMASH output were carefully examined. An operon spanning 21 kb was identified from *P. temperata* He86 containing an NRPS with two linear modules and 11 open reading frames. Although AntiSMASH did not pick up the loading domain in the first module of the NRPS, the NCBI Conserved Domain tool (13) was able to locate an acyl-CoA dehydrogenase (ACAD) in the upstream region of the first module. Therefore, the NRPS gene (gene F) in the operon contained seven identified linear modules: acyl-CoA dehydrogenase (ACAD), adenylation (A), acyl carrier protein (ACP), condensation (C), adenylation (A), acyl carrier protein (ACP), and thioesterase (TE) (Figure 2 and Table 2). The ACP module is commonly involved in loading the initial amino acid, while the TE domain is commonly involved in the final step of NRPS biosynthesis (14–16). In between the ACAD and TE there were two linear NRPS modules. This gene matched our initial search criteria for an NRPS incorporating two amino acids. The ACAD was proposed to load the initial amino acid, which was recognized by the first adenylation domain and transferred by the first ACP to the condensation domain in the second module. In the second module, the adenylation domain specifically recognized and bonded to a second amino acid, which underwent condensation with the first amino acid to form a dipeptide. The dipeptide eventually carried by the second ACP to the TE domain where the molecule got hydrolyzed and cleaved off from the NRPS protein. (Figure 2 and Table 2). A homologous BGC was also found in *P. temperata K122*, without the A, B, and C genes, suggesting the A, B and C genes were not essential in the biosynthesis of speirobactin. Further analysis with NRPSPredictor2 (17) showed weak substrate specificity prediction for both adenylation domains; the Stachelhaus sequences (18) for the two adenylation domains were identified as DYWQIGFIDK and DIWYVGACSK, both giving low matching scores for substrate specificity for tryptophan and asparagine. A tryptophan halogenase (gene D) was identified upstream to the NRPS, a carbamoyl transferase (gene G) and an aspartate racemase (gene J) was identified downstream. The isolated speirobactin contains both (6S)- and (6R)-stereoisomers, suggesting the corresponding adenylation domain incorporating both L- and D-aspartate generated by the aspartate racemase (gene J). (6R)-speirobactin utilizing D-aspartate was the more active stereoisomer, suggesting the racemase was part of the biosynthetic machinery to produce D-aspartate. Therefore, we hypothesize that the 21 kb NRPS BGC is responsible for the production of speirobactin.

**Figure 1.**
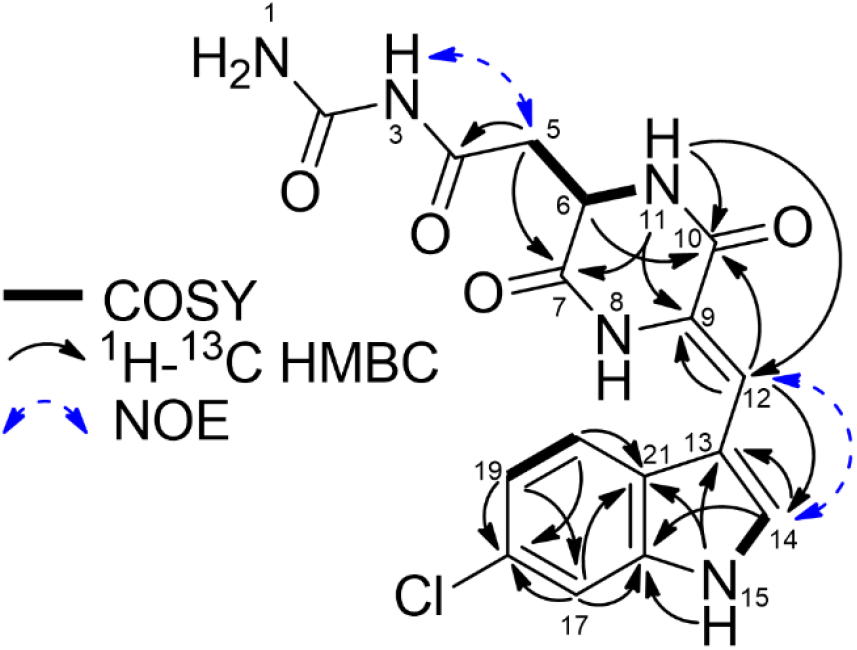
Key 2D NMR correlation of speirobactin in DMSO-*d*_*6*_.

**Table 2.** Biosynthetic genes identified from the *Photorhabdus* and *Xenorhabdus* genomes.

| Biosynthetic enzyme | Nucleotide location | GC content | Number of amino acids | Proposed function in speirobactin biosynthesis | Homologous protein from NCBI, organism, identity%/coverage%, accession No. |
| --- | --- | --- | --- | --- | --- |
| A | 289,514-290,719 | 39% | 401 | Transporter | MFS transporter, *Photorhabdus tasmaniensis*, 99/100, WP_133816297.1 |
| B | 290,986-291,420 | 33% | 144 |  | Hypothetical protein, *Photorhabdus tasmaniensis*, 100/100, WP_133816296.1 |
| C | 292,038-292,235 | 37% | 65 |  | No significant similarity found |
| D | 292,556-294,112 | 36% | 518 | Tryptophan halogenase | Tryptophan halogenase, *Photorhabdus thracensis*, 96/96, AKH65769.1 |
| E | 294,859-295,614 | 31% | 251 | Self-resistance | Pentapeptide repeat-containing protein, *Photorhabdus kayaii*, 87/100, NDL14150.1 |
| F | 295,800-303,953 | 37% | 2717 | Non-ribosomal peptide synthetase | Amino acid adenylation domain-containing protein, *Xenorhabdus* sp. 42, 76/100, MBD2820043.1 |
| G | 304,120-305,892 | 40% | 590 | Carbamoyl transferase | Carbamoyl transferase, *Photorhabdus heterorhabditis*, 91/100, KOY60149.1 |
| H | 305,921-308,053 | 39% | 710 |  | SGNH/GDSL hydrolase family protein, *Photorhabdus cinerea*, 84/100, WP_166305217.1 |
| I | 308,184-309,368 | 37% | 394 | Transporter | MFS transporter, *Xenorhabdus* sp. M, 82/99, MBD2801864.1 |
| J | 309,460-310,194 | 43% | 244 | Aspartate racemase | Aspartate/glutamate racemase family protein, *Xenorhabdus* sp. CUL, 86/96, MBD2791922.1 |
| K | 310,307-310,573 | 46% | 88 |  | Hypothetical protein, *Photorhabdus thracensis*, 97/98, AKH65770.1 |
| F1 | 1,311,013-1,318,398 | 36% | 2461 | Truncated NRPS (ACAD-A-ACP-C-A-ACP) | Hypothetical protein, *P. laumondii* subsp. *clarkei*, 80/99, RAW88993.1 |
| L | 1,318,422-1,319,624 | 50% | 400 | Transposase | IS256 family transposase, *P. luminescens*, 99/100, PQQ39455.1 |
| F2 | 1,319,722-1,320,480 | 37% | 252 | Truncated NRPS (TE) | Hypothetical protein, *P. thracensis*, 100/100, AKH62816.1 |
| M | 2,189,852-2,190,118 | 43% | 88 | Transposase | IS5 family transposase, *P. kayaii*, 80/94, NDL28030.1 |

Note: Genes A-K were from the *P. temperata* He86 circular genome, genes F1, L, and F2 were from *P. thracensis* DSM 15199, gene M was from *X. szentirmaii* DSM 16338.

**Figure 2.**
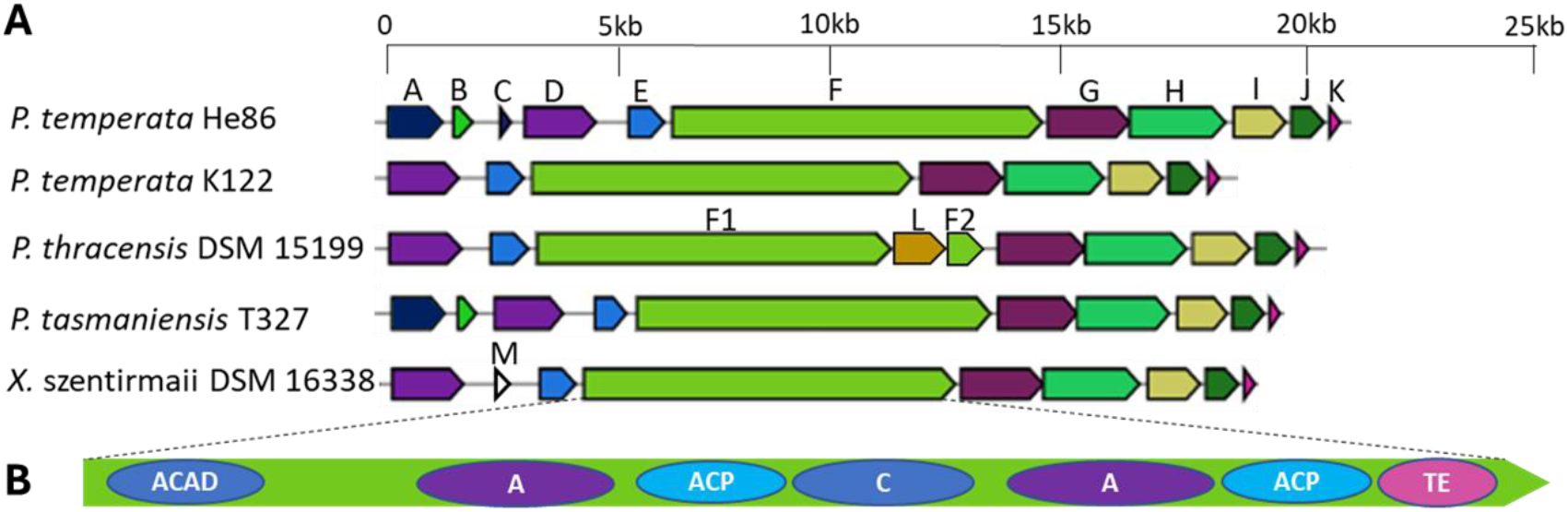
Biosynthetic gene cluster of speirobactin. (A) Alignment of biosynthetic genes across *Photorhabdus* and *Xenorhabdus* strains. (B) Domains identified in the NRPS gene in the BGC. ACAD: acyl-CoA dehydrogenase; A: adenylation; ACP: acyl carrier protein; C: condensation; D, thioesterase; E, pentapeptide repeat protein.

We also found homologous BGCs in *P. thracensis* DSM 15199, *P. tasmaniensis* T327, and *Xenorhabdus szentirmaii* DSM 16338 based on their genomes in the NCBI (Table 2). To confirm the link between the identified BGC and speirobactin, we acquired these strains and analyzed their fermentation extracts by UPLC-HRMS/MS. We were able to identify speirobactin from extracts of *P. tasmaniensis* T327 and *X. szentirmaii* DSM 16338 based on exact mass, MS^2^ pattern and retention time, but not from *P. thracensis* DSM 15199. Detailed analysis of the NRPS gene in *P. thracensis* DSM 15199 showed an insertion sequence (IS) between the ACP domain of the second module and the TE domain. The insertion sequence (gene L in Figure 2) has high homology to IS256 family transposase, an IS widespread in enterococci and staphylococci (19). We reason that the truncation of the NRPS gene in *P. thracensis* DSM 15199 by an IS resulted in the lack of speirobactin production. We identified gene M, an IS5 family transposase in the homologous BGC of *X. szentirmaii* DSM 16338 (Figure 2). The absence of genes A, B and C in the speirobactin-producing *X. szentirmaii* DSM 16338 suggests these three genes are not essential for its production. The speirobactin BGC is restricted to *Photorhabdus* and *Xenorhabdus*, based on the search of NCBI databases. However, several genes of this BGC have unusually low GC content, 31-33%, compared to 40% for the rest of the genome. This operon was probably horizontally acquired (or assembled from horizontally acquired components) from an unknown microorganism.

We noticed that the BGC of speirobactin contained a gene (E) coding for a pentapeptide repeat-containing protein (PRP) (Table S5). The best-studied member of this family is QnrA that confers resistance to fluoroquinolones. QnrA and its homologs are carried on transmissible plasmids and are present in clinical isolates of *Klebsiella pneumoniae, E. coli* and other Enterobacteriaceae. The mechanism of resistance is highly unusual, QnrA folds into a quadrilateral β-helix structure that mimics DNA which enables it to bind and sequester DNA gyrase, the target of fluoroquinolone antibiotics. Normally, diminishing the level of a target leads to increased antibiotic susceptibility. However, fluoroquinolone antibiotics act by corrupting their target rather than inhibiting its function. Fluoroquinolones stabilize the transient gyrase/DNA cleavage complex, leading to double strand brake and cell death. Diminishing the level of active gyrase will then result in partial resistance. Notably, a PRP protein, McbG was identified as an immunity factor in an *E. coli* operon coding for Microcin B17, an antimicrobial peptide targeting the DNA gyrase (20).

A Qnr homolog MfpA (mycobacterial fluoroquinolone resistance protein) was also reported in *M. tuberculosis, M. smegmatis, M. bovis, M. ulcerans*, and *M. avium* (21). The closest homologs of gene E in the speirobactin BGC were restricted to *Photorhabdus* and *Xenorhabdus* in the NCBI database. We reason that the *Photorhabdus* PRP is a self-resistance gene and the target of speirobactin is the DNA gyrase.

### Molecular Target

In order to identify the molecular target of speirobactin, *M. tuberculosis* cells were plated on a medium containing a high concentration of compound, 125 µg/mL. Resistant colonies appeared with a frequency of 1.5×10^−8^. Whole genome sequencing of four stable speirobactin resistant mutants was performed by illumina sequencing. All mutants carried mutations in the DNA gyrase; no other mutations were shared among these strains (Table 3; all mutations in Table S6). In order to validate the mutations, *gyrA* D89G and Q101H were recreated in wild-type *M. tuberculosis* by allelic replacement. Both mutations conferred resistance to speirobactin at levels over 128 µg/mL. We were not able to recreate the R128K mutation in the fresh background of the wild type.

**Table 3.** DNA gyrase mutation sites in speirobactin resistant *M. tuberculosis*.

| Mutant No. | position | Mutation | Annotation | Gene | MIC |
| --- | --- | --- | --- | --- | --- |
| 1 | 7567 | A→G | D89G(GAC→GGC) | *gyrA* | >128 $\mu\text{g/mL}$ |
| 2 | 7604 | G→C | Q101H(GAC→GGC) | *gyrA* | 32$\mu\text{g/mL}$<sup>a</sup> |
| 3 | 7567 | A→G | D89G(GAC→GGC) | *gyrA* | >128 $\mu\text{g/mL}$ |
| 4 | 7684 | G→A | R128K(AGG→AAG) | *gyrA* | >128 $\mu\text{g/mL}$ |

<sup>a</sup> MIC measurement from strain obtained in the mutant selection experiments; when the same mutation was created from wild-type Mtb by single stranded recombineering, MIC was >128 $\mu\text{g/mL}$ .

According to the crystal structure of *M. tuberculosis* GyrA (*Mt*Gyr59) (22), D89, Q101, and R128 are invariant sites close to the active site of Y129, mutations in these cites likely affect the activity of the enzyme. These results confirm GyrA as the target of speirobactin. Significantly, very high resistance of mutants shows that speirobactin does not have off-target activity.

Fluoroquinolones are the most important gyrase-targeting drugs for clearing serious *M. tuberculosis* infections in the clinic. Most bacteria have two type II topoisomerases: DNA gyrase and topoisomerase IV. Interestingly, *M. tuberculosis* only has a DNA gyrase as the sole type II topoisomerase, and it is commonly assumed to perform the function of both gyrase and topoisomerase IV, ie relaxing the supercoiling and unlinking the replicated chromosomes.

We tested the bactericidal activity of speirobactin and compared it to moxifloxacin that is used to treat tuberculosis, in exponential and stationary phase cells. Speirobactin was able to kill both exponential and stationary phase *M. tuberculosis*, though killing was somewhat weaker than moxifloxacin (Figure 3A,B).

**Figure 3.**
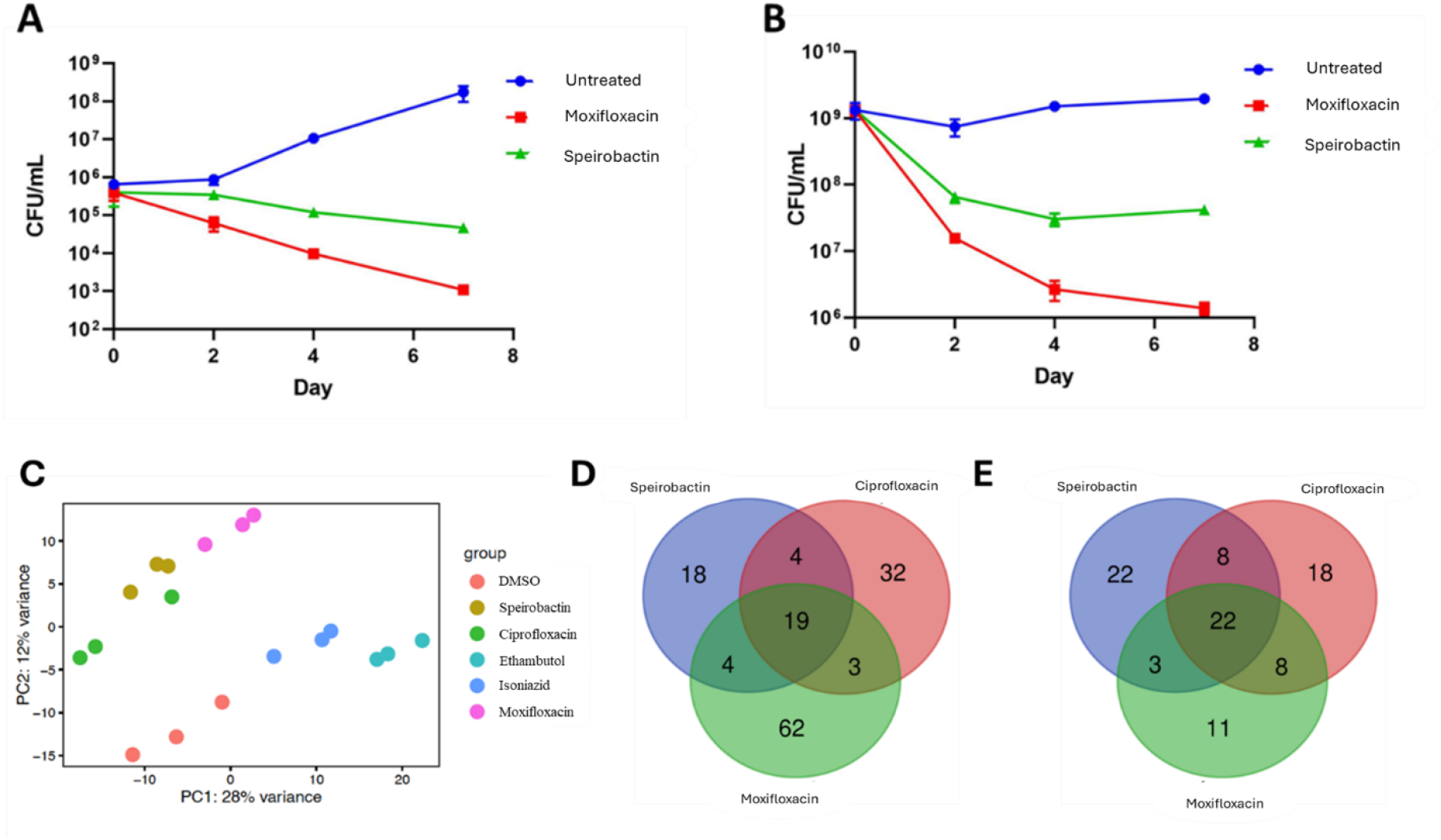
Action of speirobactin against *M. tuberculosis*. (A) Killing of *M. tuberculosis* by speirobactin and moxifloxacin in mid-exponential phase. (B) Killing of *M. tuberculosis* by speirobactin and moxifloxacin stationary phase *M. tuberculosis*. Cultures were treated with 10X MIC (MIC for both compounds is 0.125 µg/mL). The experiment was performed with biological triplicates and error bars representing standard deviations. (C) Principal component analysis of RNA-seq data where *M. tuberculosis* was treated with 2X MIC of speirobactin, moxifloxacin, ciprofloxacin, isoniazid and ethambutol for 4 hours (detailed transcriptome analysis in Supplemental File B). (D) Venn diagram for genes upregulated shared by speirobactin, moxifloxacin, and ciprofloxacin (detailed gene descriptions in Supplemental File C). (E) Venn diagram for genes downregulated shared by speirobactin, moxifloxacin, and ciprofloxacin (detailed gene descriptions in Supplemental File C).

To further understand the mode of action of speirobactin, RNA-seq analyses were performed in *M. tuberculosis* comparing speirobactin, moxifloxacin, ciprofloxacin and two front-line antibiotics isoniazid and ethambutol. While moxifloxacin and ciprofloxacin are known as DNA gyrase targeting antibiotics, isoniazid and ethambutol are cell-wall targeting. In order to prevent a complete halt in transcription but still capture compounds’ transcriptional signature, bacteria were treated at 2x MIC for a short period of 4 hours. Speirobactin clustered in between that of moxifloxacin and ciprofloxacin and away from the cell wall inhibitors and untreated samples (Figure 3C).

Based on resistance mutations, transcriptome analysis and the location of a PRP within the speirbactin BGC studies, *gyrA* is predicted to be the molecular target of speirobactin. Expression of *gyrA* (*rv0006*) was not impacted by speirobactin treatment, which is consistent with moxifloxacin and ciprofloxacin (detailed transcriptome analysis in Supplemental File B). However, there was a noted upregulation of genes involved in DNA repair that was shared between these three treatment groups (Figure 3D, Table S7, detailed gene descriptions in supplemental File C). Downregulated genes shared between speirobactin and the fluoroquinolones were involved with various metabolic processes and did not show an obvious transcriptional signature (Figure 3E, Table S8, detailed gene descriptions in Supplementary File C).

In addition to the differentially expressed genes that were shared between speirobactin and the fluoroquinolones, there were 22 downregulated genes and 18 upregulated genes that were unique to speirobactin. Analyses via STRING showed no significant enrichment detected in the upregulated genes and minor signatures related to transport (in particular the Mce2 operon) detected in the downregulated genes (Figs. S13 and S14).

We further tested the cross-resistance effects between speirobactin and moxifloxacin. Speirobactin resistant mutants were susceptible to moxifloxacin with their MIC values close to the wild-type MIC (Table 4). However, behavior of moxifloxacin resistant mutants was more complex and depended on the site of mutation: the G88C mutant was fully resistant to speirobactin while D94N was susceptible to it. These results show that although both compounds targeted DNA gyrase, their mode of action is likely to be different.

**Table 4.**
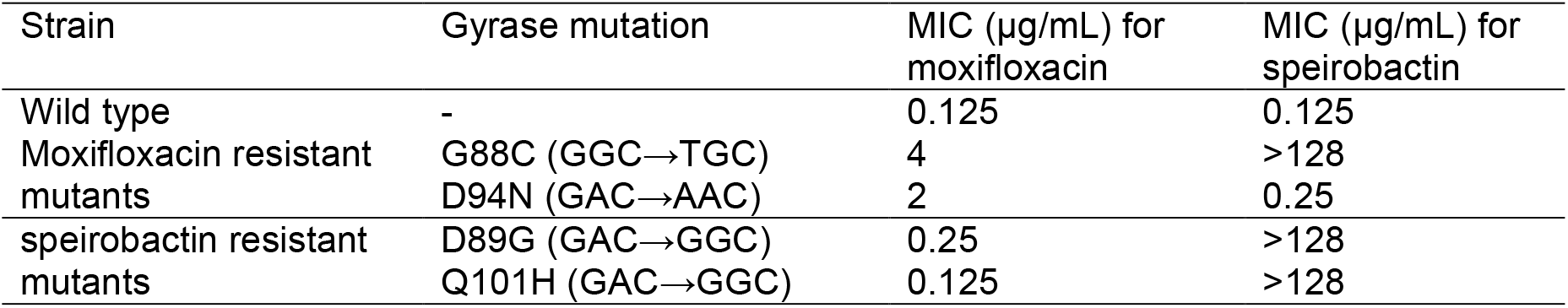
Cross-resistance analysis between moxifloxacin and speirobactin. The moxifloxacin resistant mutants and the speirobactin resistant mutants were further tested against speirobactin and moxifloxacin respectively.

## Materials and Methods

### Screening Conditions

A total of 17 *Photorhabdus* strains (Table S1) were fermented in 10 mL in 50 mL Falcon tubes using 15 nutritionally diverse media (Table S3) at 28°C and 200 rpm and 1 mL of the fermentation were sampled on day 2, day 5 and day 8. The samples were centrifuged at 8000 G and the supernatants were transferred into 96 well deep well plates, which were dried under vacuum and re-suspended in DMSO to give 1X and 15X concentrates. To perform the assay, *M. tuberculosis* [H37Rv mc2 763 6020 (ΔlysA ΔpanCD) expressing mCherry (ΔlysA ΔpanCD, pBEN_mCherry kanr 764)] cultures were diluted into fresh medium to a final OD_600_ of 0.003 and 147 µL of the diluted cultures were transferred to a 96-well black, clear-bottom microtiter plate (Corning Catalog # 07-200-567) containing 3 µL of 1X or 15X concentrated *Photorhabdus* fermentation supernatant. The assay plates were incubated for 7 days at 37°C and 100 rpm before OD_600_ measurement and fluorescence emission measurement at 610 nm with excitation at 580 nm in a plate reader. The extract was deemed active if there was over 75% growth inhibition compared to non-treatment controls.

The same extracts were tested against *S. aureus* in a disc diffusion assay on agar. Three microliters of the 1X or 15X concentrated *Photorhabdus* fermentation supernatant was spotted on an agar lawn pre-inoculated with log-phase *S. aureus* cultures. The agar plates were incubated at 37°C overnight and observed for zone of inhibition.

### Identification of speirobactin

*Photorhabdus temperata* He86 and *Photorhabdus temperata* K122 were scaled up in 4 L fermentation medium (Bacto peptone 10 g/L, K_2_HPO_4_ 1 g/L, MgSO_4_· 7H_2_O 1 g/L, (NH_4_)_2_SO_4_ 2 g/L, CaCO_3_ 2 g/L, and NaCl 10 g/L) for 5 days at 28°C and 200 rpm. The fermentations were extracted twice with equal volumes of EtOAc for two hours. The EtOAc layers were combined, dried in vacuo and fractionated with column chromatography packed with 60Å SelectraSorb Bulk Sorbent C18 (Catalog # 029070-CG). The column was equilibrated with 10 CVs (column volumes) of 5% aq. ACN containing 0.1% formic acid. The crude extracts were loaded and eluted with 5 CVs of 5%, 25%, 50% and 75% aq. ACN containing 0.1% formic acid, before washing with 10 CVs of 100% ACN containing 0.1% formic acid. Eluants were collected manually in 15 mL fractions and dried in vacuo. Each fraction was resuspended in 30 µL 25% DMSO in water and 3 µL of the suspension was transferred to a 96-well black, clear bottom microtiter plate for *M. tuberculosis* assay as described above.

The active fractions in the assay were combined and fractionated using an Agilent 1260 Affinity II HPLC system with a Restek Ultra C18 5µm 250 × 10 mm 100Å column (Catalog # 9174577). The mobile phase consisted of solvent A (H_2_O with 0.1% formic acid) and solvent B (ACN with 0.1% formic acid). The first 2 min was held at 2% solvent B, followed by a linear gradient over 20 min to 100% solvent B, and held at 100% for an additional 10 min. The flow rate was 5 mL/min. A UV detector monitor wavelengths at 220 nm, 280 nm, 310 nm and 340 nm. A fraction collector system operated in a time-based mode collecting a fraction every 0.3 min. Each fraction was dried in vacuo and resuspended in 30 µL 25% DMSO in water and 3 µL of the suspension was transferred to a 96-well black, clear bottom microtiter plate for *M. tuberculosis* assay as described above. Pure speirobactin in a fraction was identified as the active compound in the bioassay.

### Scale-up fermentation and preparative HPLC conditions

*Photorhabdus temperata* He86 was scale-up in 20 L using the aforementioned fermentation medium for 5 days at 28°C and 200 rpm. The fermentations were extracted with equal volumes of EtOAc. The EtOAc layers were combined and dried in vacuo. The EtOAc extract suspended in DMSO was further loaded onto an open column packed with SelectraSorb Bulk Sorbent C18 (Log#029070-CG) pre-conditioned with 2% aqueous ACN. Five column volumes of 2% aqueous ACN were used to remove salts and polar organic compounds. Ten column volumes of ACN were then used to elute the rest of the extract. The ACN eluent was dried in vacuo, resuspended in DMSO, then passed through a 0.2 µm filter for HPLC injections. An aliquot of 4.5 mL was injected in an Agilent 1200 series preparative HPLC system with a Primesphere C18-HC 250 mm × 50 mm, 10 µm particle size column. An isocratic condition with 25% aq. ACN with 0.1 % formic acid was used at 75 mL/min. An UV detector monitored wavelengths at 220 nm, 280 nm and 340 nm. ADC66 and speirobactin eluted at 18.5 and 23.5 min, respectively.

### Structure elucidation

Structure elucidation was conducted by 1D and 2D NMR experiments (^1^H, ^13^C, COSY, NOESY, ^1^H-^13^C HSQC, and ^1^H-^13^C HMBC) using a Bruker 900 MHz NMR, a Bruker 400 MHz NMR with a cryoprobe and a Varian 500 MHz NMR. speirobactin (2 mg) was dissolved in DMSO-*d*_*6*_ in a 5 mm NMR tube and chemical shifts were referenced to the residue solvent signal (δ_H_ 2.50, δ_C_ 39.52). LC-HRMS/MS analysis was conducted with a LTQ Orbitrap XL Hybrid Ion Trap-Orbitrap Mass Spectrometer (Thermo Scientific) in electrospray ionization positive ion mode coupled with an UltiMate 3000 RSLCnano System chromatography (Dionex). speirobactin was analyzed with a capillary column (150 mm by ID 75 µm) packed with C18 2.5mm resin (XSelect CSH C18) at a flow rate of 0.2 µL/min. The mobile phase consisted of solvent A (H_2_O with 0.1% formic acid) and solvent B (ACN with 0.1% formic acid). The first two minutes were held at 5% solvent B and then with a linear gradient increasing solvent B to 50% over 20 min.

### speirobactin

pale yellow amorphous solid; UV λmax 234, 290, 342; NMR (700 MHz, DMSO-*d*_*6*_) see Table S4; HRESIMS [M+H]^+^ *m/z* 376.0801 (calcd for C_16_H_15_N_5_O_4_Cl^+^, 376.0807, Δ = 1.60 ppm).

### Stereochemistry analysis

The stereochemistry at the aspartate moiety of speirobactin and ADC66 was analyzed with a modified Marfey’s method (23). Approximately 150 µg of speirobactin and ADC66 were hydrolyzed with 6 N HCl overnight at 100°C. The hydrolysates of the two compounds were separated into two vials for analysis with both L-FDLA and D-FDLA. The hydrolysates were dried in vacuo and then added with 1 M NaHCO_3_ (20 µL) and the choice of L-FDLA (1% solution in acetone 40 µL) or D-FDLA equivalents. Controls were prepared using L-aspartic acid reacting with both L-FDLA and D-FDLA. The reactions were maintained at 40°C for 1 h before quenching with 1 M HCl (20 µL). The samples were diluted with another 100 µL of ACN, filtered (0.2 µm) and analyzed by UPLC-HRMS/MS. An aliquot of 2 µL of each sample was injected into the Agilent Poroshell 120 EC-C18 column (3.0mm × 150mm, 2.7 µm, PN: 693975-302) with an Agilent 1260 Affinity HPLC system connected with a PDA detector and an Agilent 6530 QTOF mass spectrometer. The first two min was held at 10% solvent B (ACN with 0.1% formic acid) and 90% solvent A (H_2_O with 0.1% formic acid), then followed by a linear gradient at 65% solvent B at 27 min, followed by a linear gradient to 95% B at 27.5 min and held at 95% B until 32 min. The flow rate was 0.3 mL/min. Extracted ion chromatogram was used to monitor the reaction products.

### Chiral separation of speirobactin enantiomers

A sample in DMSO consisting of both enantiomers of speirobactin was injected onto a COSMOSIL CHiRAL Series 5C HPLC column (4.6 mm × 250 mm, Log # 15790-71) using an Agilent 1260 Infinity system monitoring UV absorbance at 342 nm. An isocratic mobile phase at 20% ACN (0.1% formic acid) and 80% H_2_O (0.1% formic acid) was used at 1 mL/min. (6R)-speirobactin eluted at 15.7 min and (6S)-speirobactin eluted at 18.9 min. The purified enantiomer was subjected to the modified Marfey’s analysis described above.

### Identification of the BGC

An overnight culture of *P. temperata* He86 and *P. temperata* K122 were used to prepare genomic DNA sample using a QIAGEN DNeasy Blood & Tissue Kit following the manufacturer’s instructions. The genomic DNA samples were for illumina sequencing by Microbial Genome Sequencing Center (MiGS; Pittsburg, PA). Nanopore sequencing was further conducted in house with a Flongle device (Oxford Nanopore Technologies) following the manufacturer’s instructions. Hybrid genome assembly using the illumina and nanopore sequencing data was conducted by Unicycler (24) using default settings. AntiSMASH (12) was used to annotate the two genomes. The homologous genes in the BGC were searched using the NCBI BlastP (13).

### Mtb growth conditions

*M. tuberculosis* MC^2^6020 (*ΔlysA ΔpanCD*) was grown in Difco 7H9 liquid media supplemented with 10% OADC, 0.5% Glycerol, 0.05% Tyloxapol, Lysine (80 µg/mL) and Pantothenate (24 µg/mL) or 7H10 solid media supplemented with 10% OADC, 0.5% Glycerol, Lysine (80 µg/mL) and Pantothenate (24 µg/mL). For screening, *M. tuberculosis* expressing a plasmid encoding the fluorescent protein mCherry was cultured in 7H9 complete media containing kanamycin and incubated at 37°C and 100 rpm. The culture was diluted into fresh medium to a final OD_600_=0.003 and aliquoted in a black clear bottom 96 well plate with bacterial extract. Final dilution of extract was 1:100. The plate was incubated for 7 days at 37°C and 100 rpm, at which point the fluorescence with excitation at 580 nm and emission at 610 nm were measured on a plate reader. The extract was deemed to have activity against *M. tuberculosis* if it had ≥75% growth inhibition when compared to the growth control. The assay was repeated for confirmation of activity.

### Minimum inhibitory concentration (MIC)

For *M. tuberculosis* assays, a final OD_600_ of 0.003 was obtained by diluting an exponentially growing culture into supplemented 7H9 medium (10% Middlebrook Oleic Albumin Dextrose Catalase (OADC) Growth Supplement (Millipore Sigma), 5% glycerol, 1% casamino acids, 0.05% tyloxapol, 80 µg ml^-1^ lysine, 24 µg ml^-1^ pantothenate. The plates were incubated 37°C and 100 rpm for 7 days. For all other microbes, a final OD_600_ of 0.001 was obtained by diluting an exponentially growing culture into BBL™ Mueller-Hinton II (Cation-Adjusted). The MIC was defined as the lowest concentration of antibiotics with no visible growth.

### Mammalian cytotoxicity

Exponentially growing NIH/3T3 mouse embryonic fibroblast (ATCC CRL-1658, in Dulbecco’s Modified Eagle’s medium supplemented with 10% bovine calf serum), and HepG2 cells (ATCC HB-8065, in Dulbecco’s Modified Eagle’s medium supplemented with 10% fetal calf serum) were seeded into a 96-well flat bottom plate. After 24 hours of incubation at 37°C, the medium was replaced with fresh medium containing two-fold serial dilution of test compounds. After 72 h of incubation at 37°C, viability was determined by adding Alamar Blue (ThermoFisher) indicator at a final concentration of 10 µg/mL and IC50 was called after 3 hours.

### Mtb time-dependent killing

Exponential culture was prepared by growing *M. tuberculosis* to mid-exponential (OD_600_∼1-1.5) then back diluting to OD_600_=0.003. For stationary phase culture, *M. tuberculosis* grew for two weeks to an OD_600_∼2. Cultures were challenged with 10x MIC of compound at 37°C. At intervals, aliquots were removed, washed once in PBS, and serially diluted and plated on 7H10 media to determine c.f.u. per mL.

### Mtb mutant generation

Mutants to speirobactin were generated by plating *M. tuberculosis* on 7H10 medium containing 10x, and 100x MIC of speirobactin. The plates were incubated at 37°C for three weeks. Four 10x mutants and four 100x mutants were picked and inoculated into 10 mL 7H9 medium and grown for 2 weeks without selection. Genome sequencing and variant calling were conducted by Microbial Genome Sequencing Center (MiGS; Pittsburg, PA).

Mutations in *gyrA* conferring resistance to 100x MIC speirobactin were recreated via single stranded recombineering as in (Table 3). Sequences of oligonucleotides used to make targeted mutations were listed in Fig. S12. Targeted mutations were confirmed via PCR and illumina sequencing.

## ACKNOWLEDGMENTS

This work was supported by the NIH grant NIH RO1AI170962. We would like to thank Dr. James Berger and Dr. Glenn Hauk for helpful discussions of the project. We acknowledge the Korea Basic Science Institute, Ochang, Korea, for collecting 900 MHz NMR data.

## ADDITIONAL FILES

Supplemental File A. Supplementary Figures and Tables.

Supplemental File B. DE Analysis from RNA seq experiments.

Supplemental File C. Upregulated and downregulated genes from RNA seq analysis.

## REFERENCES

1. Sacchettini JC, Rubin EJ, Freundlich JS. 2008. Drugs versus bugs: in pursuit of the persistent predator Mycobacterium tuberculosis. Nat Rev Microbiol 6:41–52.

2. Zumla A, Nahid P, Cole ST. 2013. Advances in the development of new tuberculosis drugs and treatment regimens. Nat Rev Drug Discov 12:388–404.

3. Khisimuzi Mdluli, Zhenkun Ma. 2007. Mycobacterium tuberculosis DNA Gyrase as a Target for Drug Discovery. IDDT 7:159–168.

4. Gavrish E, Sit CS, Cao S, Kandror O, Spoering A, Peoples A, Ling L, Fetterman A, Hughes D, Bissell A, Torrey H, Akopian T, Mueller A, Epstein S, Goldberg A, Clardy J, Lewis K. Lassomycin, a ribosomally synthesized cyclic peptide, kills mycobacterium tuberculosis by targeting the ATP-dependent protease ClpC1P1P2. Chem Biol 21:509–518.

5. Quigley J, Peoples A, Sarybaeva A, Hughes D, Ghiglieri M, Achorn C, Desrosiers A, Felix C, Liang L, Malveira S, Millett W, Nitti A, Tran B, Zullo A, Anklin C, Spoering A, Ling LL, Lewis K. 2020. Novel Antimicrobials from Uncultured Bacteria Acting against Mycobacterium tuberculosis. mBio 11:10.1128/mbio.01516-20.

6. Imai Y, Hauk G, Quigley J, Liang L, Son S, Ghiglieri M, Gates MF, Morrissette M, Shahsavari N, Niles S, Baldisseri D, Honrao C, Ma X, Guo JJ, Berger JM, Lewis K. 2022. Evybactin is a DNA gyrase inhibitor that selectively kills Mycobacterium tuberculosis. Nat Chem Biol 18:1236–1244.

7. Imai Y, Meyer KJ, Iinishi A, Favre-Godal Q, Green R, Manuse S, Caboni M, Mori M, Niles S, Ghiglieri M, Honrao C, Ma X, Guo JJ, Makriyannis A, Linares-Otoya L, Böhringer N, Wuisan ZG, Kaur H, Wu R, Mateus A, Typas A, Savitski MM, Espinoza JL, O’Rourke A, Nelson KE, Hiller S, Noinaj N, Schäberle TF, D’Onofrio A, Lewis K. 2019. A new antibiotic selectively kills Gram-negative pathogens. Nature 576:459–464.

8. Miller RD, Iinishi A, Modaresi SM, Yoo B-K, Curtis TD, Lariviere PJ, Liang L, Son S, Nicolau S, Bargabos R, Morrissette M, Gates MF, Pitt N, Jakob RP, Rath P, Maier T, Malyutin AG, Kaiser JT, Niles S, Karavas B, Ghiglieri M, Bowman SEJ, Rees DC, Hiller S, Lewis K. 2022. Computational identification of a systemic antibiotic for Gram-negative bacteria. Nat Microbiol 7:1661–1672.

9. Dossena A, Marchelli R, Pochini A. 1974. New metabolites of Aspergillus amstelodami related to the biogenesis of neoechinulin. Journal of the Chemical Society, Chemical Communications 771–772.

10. Gao H, Liu W, Zhu T, Mo X, Mándi A, Kurtán T, Li J, Ai J, Gu Q, Li D. 2012. Diketopiperazine alkaloids from a mangrove rhizosphere soil derived fungus Aspergillus effuses H1-1. Organic & Biomolecular Chemistry 10:9501–9506.

11. Ling LL, Xian J, Ali S, Geng B, Fan J, Mills DM, Arvanites AC, Orgueira H, Ashwell MA, Carmel G, Xiang Y, Moir DT. 2004. Identification and Characterization of Inhibitors of Bacterial Enoyl-Acyl Carrier Protein Reductase. Antimicrob Agents Chemother 48:1541– 1547.

12. Blin K, Shaw S, Augustijn HE, Reitz ZL, Biermann F, Alanjary M, Fetter A, Terlouw BR, Metcalf WW, Helfrich EJ. 2023. antiSMASH 7.0: new and improved predictions for detection, regulation, chemical structures and visualisation. Nucleic acids research 51:W46–W50.

13. Altschul SF, Madden TL, Schäffer AA, Zhang J, Zhang Z, Miller W, Lipman DJ. 1997. Gapped BLAST and PSI-BLAST: a new generation of protein database search programs. Nucleic acids research 25:3389–3402.

14. Matsuda K. 2025. Macrocyclizing-thioesterases in bacterial non-ribosomal peptide biosynthesis. J Nat Med 79:1–14.

15. Pourmasoumi F, De S, Peng H, Trottmann F, Hertweck C, Kries H. 2022. Proof-Reading Thioesterase Boosts Activity of Engineered Nonribosomal Peptide Synthetase. ACS Chem Biol 17:2382–2388.

16. Miller BR, Gulick AM. 2016. Structural Biology of Non-Ribosomal Peptide Synthetases. Methods Mol Biol 1401:3–29.

17. Röttig M, Medema MH, Blin K, Weber T, Rausch C, Kohlbacher O. 2011. NRPSpredictor2—a web server for predicting NRPS adenylation domain specificity. Nucleic Acids Res 39:W362–W367.

18. Stachelhaus T, Mootz HD, Marahiel MA. 1999. The specificity-conferring code of adenylation domains in nonribosomal peptide synthetases. Chem Biol 6:493–505.

19. Hennig S, Ziebuhr W. 2010. Characterization of the Transposase Encoded by IS 256, the Prototype of a Major Family of Bacterial Insertion Sequence Elements. J Bacteriol 192:4153–4163.

20. Collin F, Maxwell A. 2019. The microbial toxin microcin B17: prospects for the development of new antibacterial agents. Journal of molecular biology 431:3400–3426.

21. Vetting MW, Hegde SS, Fajardo JE, Fiser A, Roderick SL, Takiff HE, Blanchard JS. 2006. Pentapeptide Repeat Proteins. Biochemistry 45:1–10.

22. Tretter EM, Schoeffler AJ, Weisfield SR, Berger JM. 2010. Crystal structure of the DNA Gyrase GyrA N-terminal domain from Mycobacterium tuberculosis. Proteins 78:492–495.

23. Vijayasarathy S, Prasad P, Fremlin LJ, Ratnayake R, Salim AA, Khalil Z, Capon RJ. 2016. C_3_ and 2D C_3_ Marfey’s Methods for Amino Acid Analysis in Natural Products. J Nat Prod 79:421–427.

24. Wick RR, Judd LM, Gorrie CL, Holt KE. 2017. Unicycler: resolving bacterial genome assemblies from short and long sequencing reads. PLoS computational biology 13:e1005595.

